# Inferring putative ancient whole genome duplications in the 1000 Plants (1KP) initiative: access to gene family phylogenies and age distributions

**DOI:** 10.1101/735076

**Authors:** Zheng Li, Michael S Barker

## Abstract

Polyploidy or whole genome duplications (WGDs) repeatedly occurred during green plant evolution. To examine the evolutionary history of green plants in a phylogenomic framework, the 1KP project sequenced over 1000 transcriptomes across the Viridiplantae. The 1KP project provided a unique opportunity to study the distribution and occurrence of WGDs across the green plants. In the 1KP capstone analyses, we used a total evidence approach that combined inferences of WGDs from Ks and phylogenomic methods to infer and place ancient WGDs. Overall, 244 putative ancient WGDs were inferred across the Viridiplantae. Here, we describe these analyses and evaluate the consistency of the WGD inferences by comparing them to evidence from published syntenic analyses of plant genome assemblies. We find that our inferences are consistent with whole genome synteny analyses and our total evidence approach may minimize the false positive rate throughout the data set. Given these resources will be useful for many future analyses on gene and genome evolution in green plants, we release 383,679 nuclear gene family phylogenies and 2,306 gene age distribution (Ks) plots from the 1KP capstone paper.

## 1. Methods

The 1000 plants (1KP) project [1] sequenced the transcriptomes of 1,173 plant species from across the green plant phylogeny. These newly sequenced data provided crucial new genomic data for previously under- or unsampled lineages of green plants. One of the major discoveries of the early era of plant genome sequence was the observation of ancient whole genome duplications (WGDs) or paleopolyploidy in the history of most sequenced plant genomes [2,3]. Despite progress on understanding the distribution of WGDs across the phylogeny of green plants, many lineages have remained unstudied for lack of data. The expansive phylogenetic sampling of the 1KP provided an opportunity to infer putative WGDs and assess their frequency and distribution across the green plant tree of life. To survey potential WGDs, we used a total evidence approach to infer and place putative ancient WGDs in the 1KP capstone phylogeny. WGDs were inferred from age distributions of gene duplications by analyzing transcriptomes of single species with the DupPipe pipeline [4]. To place inferred WGDs from Ks plots onto the species phylogeny, we compared the median paralog divergence (*Ks*) of putative WGD peaks to the divergence of orthologs among species across the phylogeny [4]. We also employed phylogenomic analyses and simulations of WGDs using MultitAxon Paleopolyploidy Search(MAPS) [5,6] to corroborate the inferences and phylogenetic placements of the putative ancient WGDs. Here we provide details of our analyses as well as Ks plots that represent each major lineages and two walkthrough examples from our 1KP capstone analyses to demonstrate our total evidence approach. Finally, we evaluate our inferences of WGDs by comparing them with evidence from published syntenic analyses of plant genome assemblies.

### 1.1 DupPipe analyses of WGDs from transcriptomes of single species

For each transcriptome, we used the DupPipe pipeline to construct gene families and estimate the age of gene duplications [4]. We translated DNA sequences and identified reading frames by comparing the Genewise [7] alignment to the best-hit protein from a collection of proteins from 25 plant genomes from Phytozome [8]. For each analysis, we used protein-guided DNA alignments to align our nucleic acid sequences while maintaining reading frame. We estimated synonymous divergence (*Ks*) using PAML [9] with the F3X4 model for each node in the gene family phylogenies. A recent study has shown that estimating the node Ks values for duplicates from gene family trees rather than pairwise comparisons of paralogs can reduce error in estimating Ks values of duplication events and has a significant impact on the resolution of WGD peaks [10]. In this project, we used the approach described in Tiley et al. 2018. Previous analyses also indicate that there is reasonable power to infer WGDs in Ks plots when paralog divergences are Ks < 2. Saturation and other errors accumulate at paralog divergences of Ks > 2 and can create false signals of WGDs and make distinguishing true WGDs from the background a fraught task [10,11]. We followed the recommendations of these studies in all of our 1KP Ks plot inferences. Although we plotted and presented two sets of histograms with x-axis scales of Ks = 2 and Ks = 5 to assess WGDs at different resolutions (Fig. 1, Fig. 2), we did not identify peaks with Ks > 2 as potential WGDs without other data available (e.g., synteny or phylogenomic evidence). Note that this means the rate of substitution in a lineage limits the depth of time at which we can reliably infer the presence or absence of putative WGDs. The 2,306 Ks plots generated in these analyses are available here: https://bitbucket.org/barkerlab/1kp/src/master/.

**Fig. 1.**
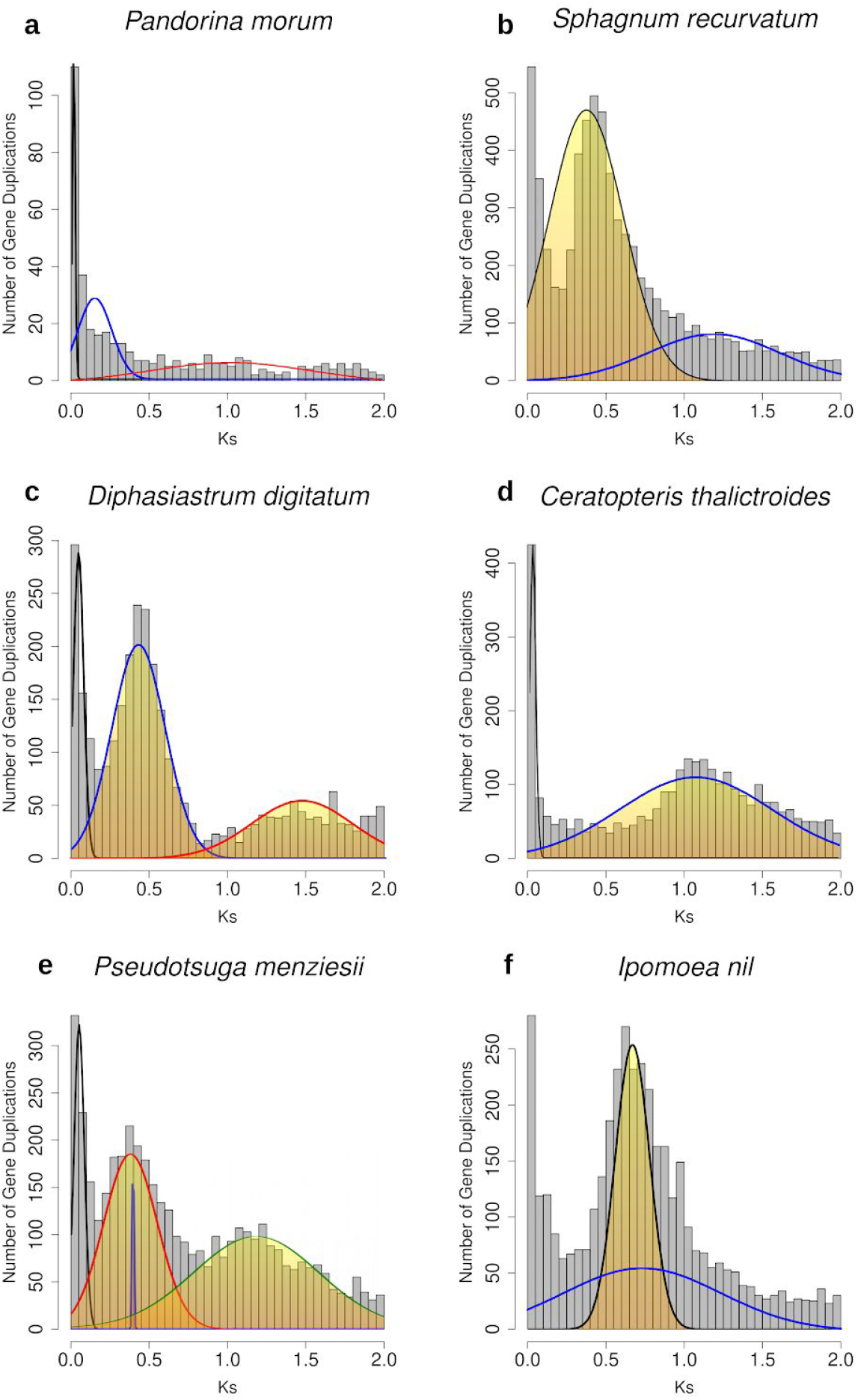
Histograms of the age distribution of gene duplications (Ks plots) with mixture models of inferred WGDs for **(a)** *Pandorina morum* (green algae), no inferred WGD peak. **(b)** *Sphagnum recurvatum* (Moss), inferred WGD peak median Ks=0.38. **(c)** *Diphasiastrum digitatum* (Lycophyte), inferred WGD peaks median Ks=0.42, 1.62. **(d)** *Ceratopteris thalictroides* (Fern), inferred WGD peak median Ks=1.08. **(e)** *Pseudotsuga menziesii* (Gymnosperm), inferred WGD peak median Ks=0.38, 1.18. **(f)** *Ipomoea nil* (Angiosperm) inferred WGD peak median Ks=0.66. Histogram x-axis scale is Ks 0–2. The mixture model distributions consistent with inferred ancient WGDs are highlighted in yellow.

**Fig. 2.**
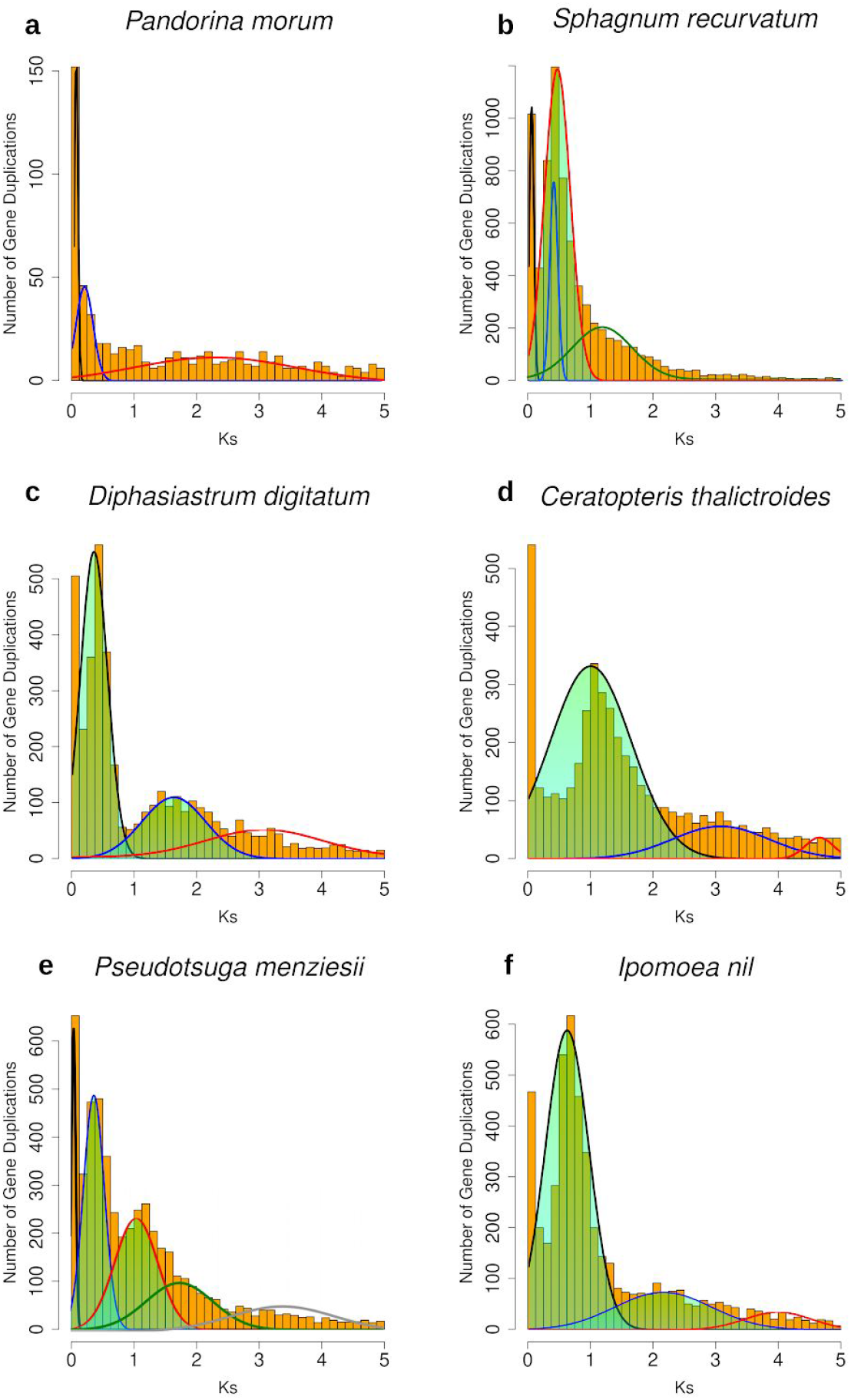
Histograms of the age distribution of gene duplications (Ks plots) with mixture models of inferred WGDs for **(a)** *Pandorina morum* (green algae), no inferred WGD peak. **(b)** *Sphagnum recurvatum* (Moss), inferred WGD peak median Ks=0.38. **(c)** *Diphasiastrum digitatum* (Lycophyte), inferred WGD peaks median Ks=0.42, 1.62. **(d)** *Ceratopteris thalictroides* (Fern), inferred WGD peak median Ks=1.08, 3.07. **(e)** *Pseudotsuga menziesii* (Gymnosperm), inferred WGD peak median Ks=0.38, 1.18. **(f)** *Ipomoea nil* (Angiosperm) inferred WGD peak median Ks=0.66, 2.15. Histogram x-axis scale is Ks 0–5. The mixture model distributions consistent with inferred ancient WGDs are highlighted in green.

To identify significant features in the gene age distributions that may correspond to WGDs, we used two statistical tests: Kolmogorov–Smirnov goodness of fit tests and mixture models. We first identified taxa with potential WGDs by comparing their paralog ages to a simulated null distribution without ancient WGDs using a K–S goodness of fit test [12]. For taxa with evidence for a significant peak relative to the null, we then used a mixture model implemented in the mixtools R package [13] to identify significant peaks of gene duplication consistent with WGDs and estimate their median Ks values (Fig. 1, Fig. 2). These approaches have been used to infer WGDs in Ks plots in many species that were subsequently corroborated by syntenic analyses of whole genome sequences [12,14–16]. There is a recent trend in the community of authors simply surveying the Ks plots of single species without a model or statistical inference to infer a WGD (e.g., [17–20]). By using these two statistical tests, our results have been more rigorously evaluated than many recent studies of WGDs.

To visually demonstrate our gene age distribution approach, we provide example Ks plots for four major lineages across the green plant phylogeny. In the green alga *Pandorina morum*, the K-S test indicated that the paralog age distribution was significantly different than a simulated null. However, we do not observe any peaks of duplication consistent with the expected signature of an ancient WGD from the two sets of histograms (Fig. 1a, Fig. 2a). In other land plant examples, the K-S test also found paralog age distributions were significantly different than null simulations (*p* = 0). In the bryophyte and fern examples, we observed single peaks of duplication consistent with an ancient WGD in the Ks plots of each species (*Sphagnum recurvatum*, median Ks = 0.3814, Fig. 1b & 2b; *Ceratopteris thalictroides*, median Ks = 1.0793, Fig. 1d & 2d). In the lycophyte, gymnosperm, and angiosperm examples, we observed two peaks of duplication consistent with two rounds of putative ancient WGD in each species. The mixtools mixture models estimated that these putative WGD peaks have median Ks of 0.4247 and 1.6229 in *Diphasiastrum digitatum* (Fig. 1c, Fig. 2c), median Ks values of 0.3724 and 1.1572 in *Pinus radiata* (Fig. 1d, Fig. 2d), and median Ks values of 0.6646 and 2.1532 in *Ipomoea nil* (Fig. 1e, Fig. 2e).

### 1.2 Estimating orthologous divergence

To place putative WGDs in the context of lineage divergence, we estimated the synonymous divergence of orthologs among pairs of species that may bracket the phylogenetic position of a WGD in our sampled taxa. Orthologs were identified as reciprocal best blast hits in pairs of transcriptomes using the RBH Ortholog pipeline [4]. This pipeline uses protein-guided DNA alignments to align our nucleic acid sequences while maintaining reading frame. The pairwise synonymous (*Ks*) divergence for each pair of orthologs is then estimated using PAML with the F3X4 model [9]. The mean and median ortholog synonymous divergences were recorded and compared to the synonymous divergence of inferred paleopolyploid peaks estimated by the mixture model. If the median synonymous divergence of WGD paralogs was younger than the median synonymous divergence of orthologs, WGDs were interpreted to have occurred after lineage divergence. Similarly, if the synonymous divergence of WGD paralogs was older than the ortholog synonymous divergence, then we interpreted those WGDs as shared by those taxa. By comparing paralog and ortholog synonymous divergences, we placed inferred ancient WGDs in a phylogenetic context. To better demonstrate this ortholog divergence analysis, we provide a walk through example using a putative WGD inferred in the ancestry of the Pinaceae in section 2.

### 1.3 MAPS analyses of WGDs from transcriptomes of multiple species

We used MAPS, a gene tree topology sorting algorithm [5,6], to confirm the placement of ancient WGDs that may be shared by at least three species. MAPS uses a given species tree to filter collections of nuclear gene trees for subtrees consistent with relationships at each node in the species tree. For each MAPS analysis, gene families were clustered using OrthoFinder [21] with reciprocal protein BLAST (blastp) searches using an E-value of 10e-5 as a cutoff. Gene families were clustered using the default parameters of OrthoFinder. We filtered the gene family clusters to include only gene families that contained at least one gene copy from each taxon. We constructed alignments and phylogenies for each gene family using PASTA [22]. For each gene family phylogeny, we ran PASTA until we reached three iterations without an improvement in likelihood score using a centroid breaking strategy. Within each iteration of PASTA, we constructed subset alignments using MAFFT [23], employed Muscle [24] for merging these subset alignments, and RAxML [25] for tree estimation. The parameters for each software package were the default options for PASTA. We used the best scoring PASTA tree for each multi-species nuclear gene family to collectively estimate the numbers of shared gene duplications on each branch of the given species. To maintain sufficient gene tree numbers to infer ancient WGDs, we used collections of gene trees for six to eight taxa for each MAPS analysis. The entire collection of 383,679 nuclear gene family phylogenies generated for all MAPS analyses is available here: https://bitbucket.org/barkerlab/1kp/src/master/.

We selected taxa for our MAPS analyses to minimize potential mapping errors at the tips and roots of species trees. Gene tree error may create a bias that causes more gene losses to map at the tips and more gene duplications to map to roots in gene tree reconciliation analyses [26]. Although there is not a general solution to this problem, we used two different approaches in our MAPS analyses to minimize the impact of this known issue. First, we expect the tips and roots of our MAPS analyses to have much higher duplication mapping error. Given that the numbers of subtrees at the tips and roots may be skewed, we have lower confidence in estimates at the tip and root nodes compared to the number of mapped duplications in the center of our MAPS phylogenies. For this reason, we aimed to place the focal WGD test node in the middle of the phylogeny being examined. Secondly, we implemented an option in MAPS to increase taxon occupancy in the gene trees by requiring a minimum number of ingroup taxa be present in each subtree [5]. Based on previous work [27] and balancing the number of trees retained in our analyses, we used a minimum 45% ingroup taxa requirement in our MAPS analyses. If this minimum ingroup taxa number requirement is not met for a gene tree, it will be filtered out and excluded from our analysis. As we discussed in Li et al. 2018, requiring higher taxon occupancy greatly reduced the bias of mapping duplications to older nodes of the phylogeny as observed by Hahn (2007) and led to less inflated estimates of duplications on deeper nodes (Fig. 3).

**Fig. 3.**
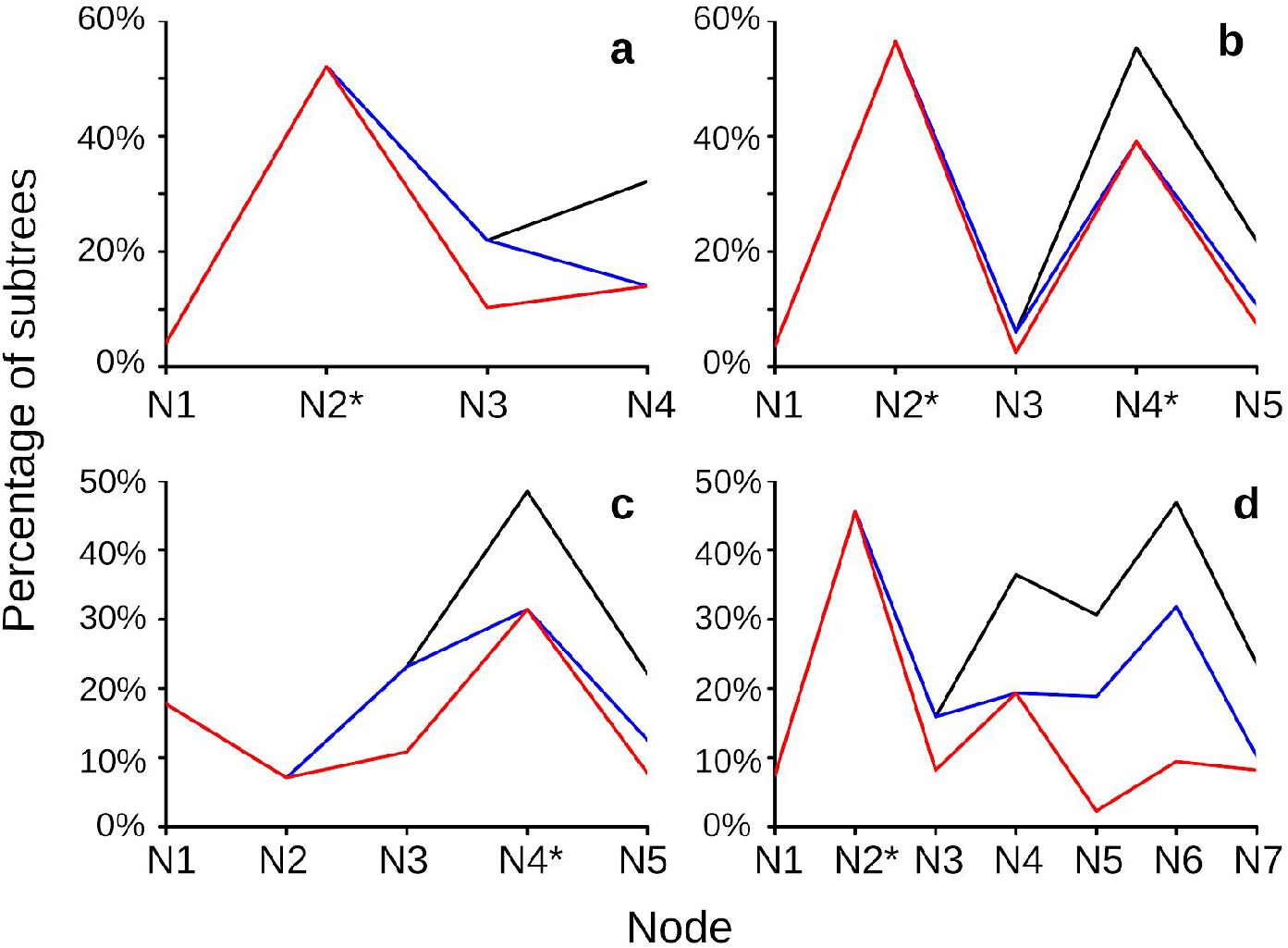
Increasing taxon occupancy decreases the inflation of mapped duplications towards the root of the species tree in MAPS. The black line represents the MAPS result without the minimum taxon requirement. The blue line represents the MAPS results with a 35% minimum taxa requirement. The red line represents the MAPS results with a 45% minimum taxa requirement. N1 corresponds to the tip node, and the last node (eg: N4 in **(a)**) corresponds to the root node. * represents nodes associated with inferred WGDs. **(a)** 1KP MAPS result of eudicot ancient hexaploidy event, N2 represents the node associated with this paleohexaploidy event. See MAPS E21 in the One Thousand Plant Transcriptomes Initiative, 2019 for details. **(b)** N2 represents the node associated with an inferred Pinaceae WGD, N4 represents the node associated with the inferred seed plant WGD. See Fig. 4c, d for the phylogeny. **(c)** N4 represents node associated with the paleohexaploidy event shared by most Compositae, see Fig. 5a for the phylogeny. **(d)** N4 represents node associated with the Heliantheae ancient WGD, see Fig. 5b for the phylogeny.

As genomic data has expanded, methods for inferring WGDs from phylogenetic analyses have matured over time to include more formal approaches for assessing WGDs. The increased taxon sampling present in larger datasets has allowed the field to begin analyzing genomic data from multiple related species that may have a shared WGD in their ancestry. Some early phylogenomic approaches simply used a hard cutoff based on numbers or percentages of gene trees to label an episode of gene duplication a putative WGD [18]. Although many WGDs may be inferred because of large changes in duplication numbers across a phylogeny, gene duplications vary across the phylogeny because of changes in branch length and variation in gene birth and death rates. We introduced simulations and statistical analyses in MAPS to address some of the issues associated with the phylogenomic inference of ancient WGDs [5]. Ancient WGDs are inferred in two steps in the MAPS framework. We first develop a null simulation of the number of expected duplications on each branch of our species tree based on a range of estimated background gene birth and death rates. The null simulation used gene birth and death rates estimated from each tree using WGDgc as described in [28], and used the GuestTreeGen program from GenPhyloData [29] to generate simulated gene trees as described in Li et al. (2018). This null simulation accounts for variation in the number and percent of gene duplications associated with branch length and background birth/death rates among the sampled taxa. Significant bursts above this null indicate a deviation from the background birth and death rate as expected for episodic events like WGDs. We used Fisher’s exact test to compare our observed MAPS results to the null simulations and identify significant episodes of duplication. All nodes are compared against the null model to identify significant episodes of gene duplication across a species tree. Once these significant episodes of gene duplication are identified, we used a second set of gene tree simulations to assess if they were consistent with a WGD. Again, we used Fisher’s exact test to compare our observed numbers of duplications to the number of shared duplications expected with a WGD at a particular location in the phylogeny. If these increases in gene duplications were caused by a WGD, then we expect the numbers of shared gene duplications among extant taxa to be consistent with these positive simulations. By using these simulations and statistical methods, MAPS explicitly accounts for the number of duplications expected on branches of different lengths within species trees and provides a statistical test to assess if an episode of duplication is consistent with a potential ancient WGD.

It should be emphasized that we used a total evidence approach to infer WGDs in the 1KP capstone project. We combined evidence from single species Ks plots, pairwise ortholog divergence analyses, and multispecies MAPS analyses to identify ancient episodes of gene duplication consistent with WGDs and place them on our species tree. For example, we did not call a WGD based only on evidence from a MAPS analysis. In the few cases where the results of our different inference approaches conflicted, we relied on the weight of evidence from a majority of analyses and, if available, other analyses from the literature to infer a putative WGD. These were mostly cases where inferences from Ks plots, ortholog comparisons, and the previous literature agreed, but MAPS did not. In these cases, we recognized the event as a significant burst of gene duplication and indicated this in Supplemental text and tables, and labeled as blue squares on the ED WGD Phylogeny Figure [1]. These events may be WGDs that should be analyzed in subsequent analyses with new data or methods.

## 2. Walk-through examples

To better demonstrate our approach for inferring ancient WGDs, we selected two examples from the 1KP analyses as walk-throughs. We chose the Pinaceae and Compositae ancient WGD analyses as examples (Fig. 4, 5) because these analyses represent different scales and complexities of duplication events. Previous analyses have found evidence for two rounds of WGD in the history of the Pinaceae [6], including a potential WGD in the ancestry of all seed plants [6,30]. However, other analyses have questioned the placement and/or existence of significant bursts of gene duplication in these lineages [31,32]. In contrast, the Compositae walk-through example has no conflict among studies, but is a complex nested paleohexaploidy in the ancestry of one of the largest families of flowering plants [33,34]. Inferring the location of the nested WGDs that comprise the paleohexaploidy, while also distinguishing other WGDs in these data, is a potentially challenging task for transcriptome based phylogenomic analyses. Below, we walk through our results for these examples and explain how we arrived at our inferences of a WGD (or not). It should be noted that we conducted a similar level of analysis and decision making process in the inference of all 244 putative WGDs in the 1KP capstone analysis.

**Fig. 4.**
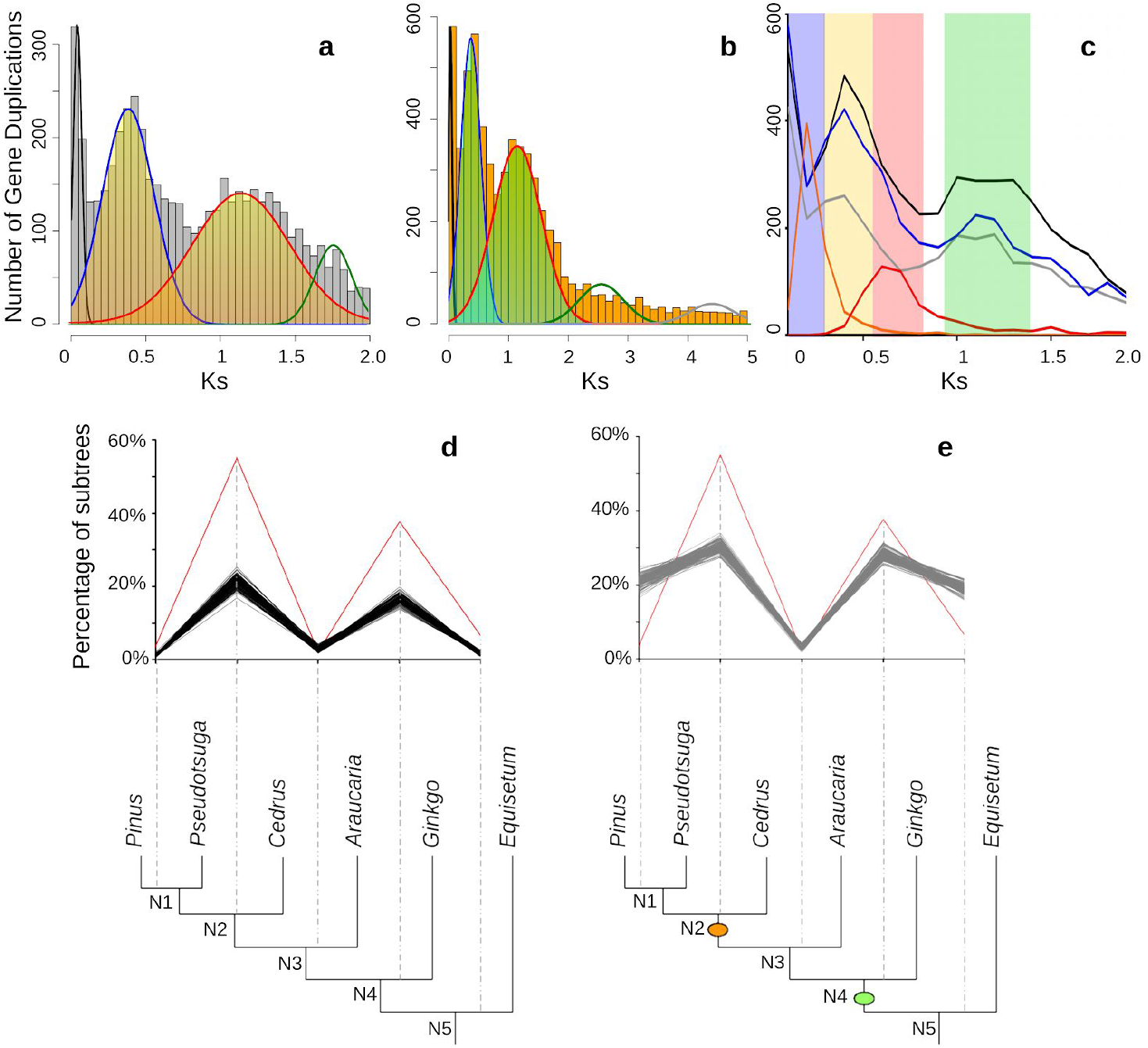
Histograms of the age distribution of gene duplications (Ks plots) with mixture models of inferred WGDs for *Pseudotsuga menziesii* (Gymnosperm), inferred WGD peak median Ks=0.37, 1.16. **(a)** Histogram x-axis scale is Ks 0–2. The mixture model distributions consistent with inferred ancient WGDs are highlighted in yellow. **(b)** Histogram x-axis scale is Ks 0–5. The mixture model distributions consistent with inferred ancient WGDs are highlighted in green. **(c)** Combined Ks plot of the gene age distributions of *P. menziesii* (blue), *Pinus radiata* (black), *Cedrus libani* (gray), and ortholog divergences of *Pinus* vs. *Cedrus* (orange) and *Cedrus* (Pinaceae) vs. *Cephalotaxus* (Cephalotaxaceae) (red). The median peaks for these plots are highlighted. **(d)** and **(e)** MAPS results from observed data, null and positive simulations on the associated phylogeny. **(d)** Percentage of subtrees that contain a gene duplication shared by descendant species at each node, results from observed data (red line), 100 resampled sets of null simulations (multiple black lines). **(e)** Percentage of subtrees that contain a gene duplication shared by descendant species at each node, results from observed data (red line), and positive simulations (multiple gray lines). The orange oval corresponds to the location of an inferred WGD in Pinaceae. The green oval corresponds to the location of an inferred WGD in seed plants.

**Fig. 5.**
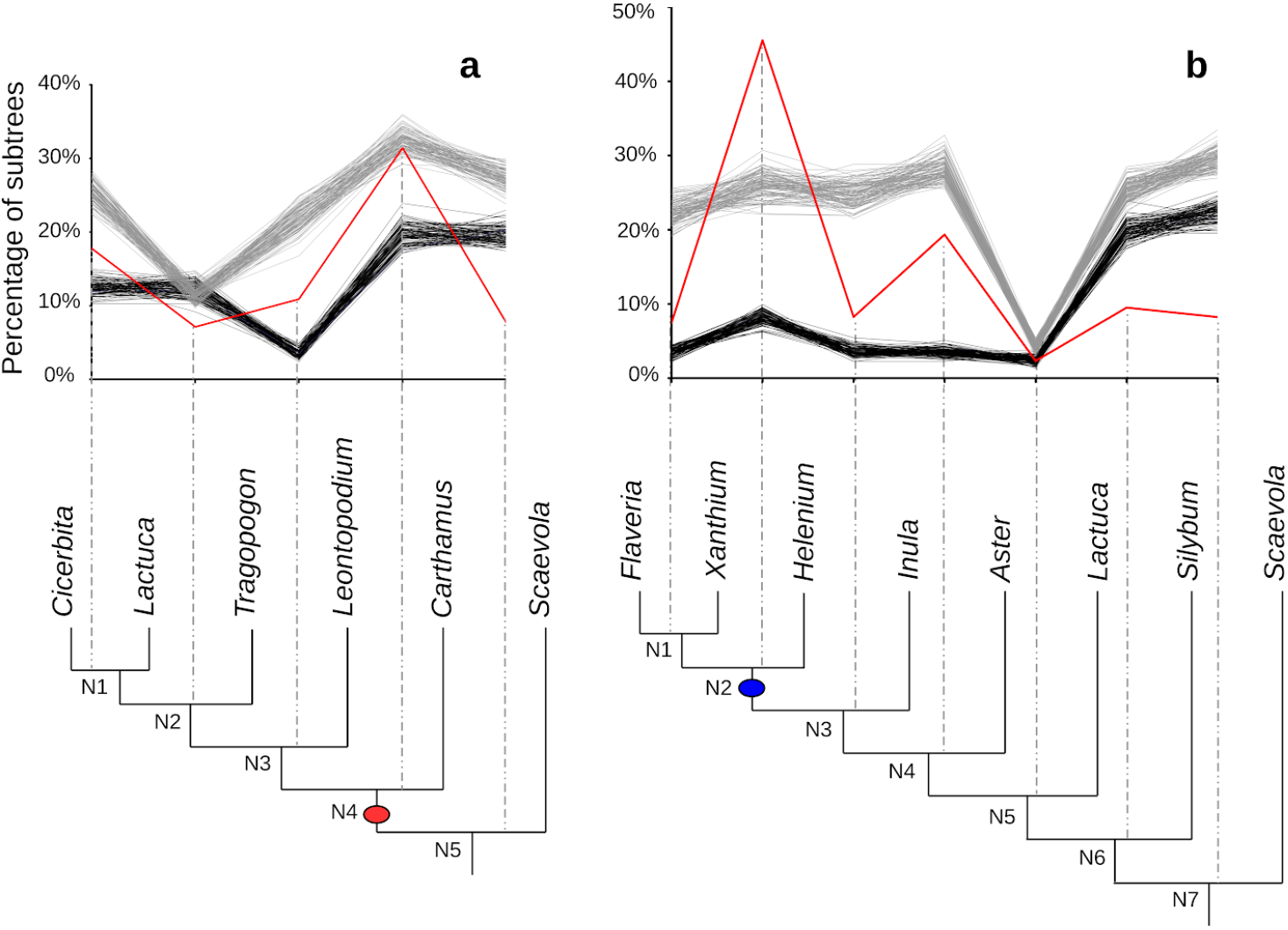
Asteraceae MAPS results from observed data, null, and positive simulations on the associated phylogeny. **(a)** Percentage of subtrees that contain a gene duplication shared by descendant species at each node, results from observed data (red line), 100 resampled sets of null simulations (multiple black lines) and positive simulations (multiple gray lines). The red oval corresponds to the paleohexaploidy event in the Compositae. **(b)** Percentage of subtrees that contain a gene duplication shared by descendant species at each node, results from observed data (red line), 100 resampled sets of null simulations (multiple black lines) and positive simulations (multiple gray lines). The blue oval corresponds to the Heliantheae ancient WGD.

Consistent with previous research [6,30], we observed evidence for at least two rounds of duplication in the ancestry of the Pinaceae. We observed two peaks of duplication consistent with two rounds of ancient WGDs in the history of three Pinaceae genera (*Pinus, Pseudotsuga*, and *Cedrus;* Fig. 4). Recent peaks of duplication in species of *Pinus*, *Pseudotsuga*, and *Cedrus* have a median Ks ~0.3 (Fig. 4a-c), older than their ortholog divergences (Ks ~ 0.18; Fig. 4c). These ortholog divergence analyses suggest the younger putative WGD in the three species is most likely shared by all Pinaceae. However, this putative WGD is not likely shared by other conifers because the ortholog divergences of the Pinaceae to other conifers is nearly twice the paralog divergence of the putative WGD. For example, ortholog divergences of members of the Pinaceae relative to members of the Cephalotaxaceae is Ks ~0.6 (Fig. 4c), consistent with this duplication event occurring after the divergence of these conifer families. The older peaks observed in *Pinus, Pseudotsuga*, and *Cedrus* have a median Ks ~1 (Fig. 4c), most likely shared by all seed plants but more recent than the divergence of seed plants and ferns (Ks ~3, estimated in 1KP capstone project).

As described above, we further assessed the nature of phylogenetic position of these putative WGDs using MAPS. We selected species of *Pinus, Pseudotsuga*, and *Cedrus* to represent Pinaceae in this MAPS analyses. We also selected species of *Araucaria* and *Ginkgo* to represent other gymnosperms, and species of *Equisetum* and *Selaginella* were used as outgroups. For the null simulations, we first simulated 3000 gene trees using the mean background gene duplication rate (λ) and gene loss rate (μ). We then randomly resampled 1000 trees without replacement from the total pool of gene trees 100 times to provide a measure of uncertainty of the percentage of subtrees at each node (Fig. 4d). At nodes corresponding to N1, N2, N4, and N5, we observed significantly more shared duplications than expected compared to the null simulations (*p* < 0.01) (Fig. 4d). For positive simulations, we incorporated a WGD at nodes N1, N2, N4, and N5 and simulated gene trees using the same methods described above. At the node representing the MRCA of Pinaceae (N2) and the node representing the MRCA of gymnosperms (N4), we interpreted an episodic burst of shared gene duplication that is statistically consistent with our positive simulations of WGDs (Fig. 4e). The results from our comparison to the null and positive simulations are consistent with those from Ks plots and ortholog divergence analyses described above, as well as those of our previous study in gymnosperms [6]. These results and another MAPS analysis in the 1KP capstone project (MAPS D1) show evidence consistent with a putative ancient WGD shared among all Pinaceae and another putative WGD that likely occurred in the ancestry of seed plants [1].

In addition to our analyses with the 1KP capstone dataset, other analyses have also inferred a putative WGD in the ancestry of all seed plants [6,30,31] and in the ancestry of different conifer families [6]. Consistent with our previous analyses [6], the relatively dense phylogenetic sampling of the 1KP allowed us to confirm that the putative seed plant WGD is not shared with monilophytes. A recent study proposed that cycads and *Ginkgo* might have shared another round of ancient WGD(s) [19]. However, other analyses in the 1KP capstone (MAPS D1 and related ortholog divergence analyses) using three species of cycads, *Ginkgo, Amborella*, and outgroups rejects this hypothesis. Instead, we find evidence that the signature detected by Roodt et al. (2017) in cycads and *Ginkgo* is most likely the putative seed plant WGD (One Thousand Plant Transcriptomes Initiative, 2019). In the 1KP and previous research [6], we also found evidence for other putative ancient WGDs in the ancestry of some families of conifers, including the Pinaceae as described above. Using whole genome data from *Ginkgo biloba, Picea abies*, and *Pinus taeda*, a recent study does not find evidence in both Ks plots and phylogenomic analyses for the Pinaceae WGD [31]. The absence of a putative Pinaceae WGD peak in their Ks plot is possibly due to the quality of the genome assembly and annotation, or the scaling of their Ks plot which may obscure the peaks we observed in all Pinaceae taxa. In the 1KP capstone project, we consistently observed two peaks of gene duplication consistent with putative WGDs in all Ks plots from the 14 species of Pinaceae analyzed. Only one conifer species, *Picea abies*, was included in the analysis by Zwaenepoel and Van de Peer (2019). It is possible the lack of support for the Pinaceae WGD is due to the limited sampling of conifers, as they [31] demonstrated that taxon sampling can have a significant impact on WGD inference with taxon-dependent support for the well established eudicot hexaploidy [16,35–38]. Given the evidence from Ks plots, ortholog divergence, and MAPS analyses that we discussed above, our current inference and placement of the putative Pinaceae and seed plants WGDs is currently the best explanation for these large scale gene duplication events. Future studies with new data, especially with higher quality gymnosperm genome assemblies, are needed to test these hypothesized WGDs.

To further demonstrate our total evidence approach to resolve complex ancient WGDs, we provide a walk-through of our analyses of ancient WGDs in the Asteraceae. We previously inferred two rounds of ancient WGD consistent with a paleohexaploidy in the ancestry of the Compositae [14,33,34]. The paleohexaploid nature of this WGD was later supported by synteny analyses of the sunflower and other Compositae genomes [39–41]. Given the great phylogenetic depth of sampling in the 1KP project and our introduction of a new statistical test for inferring WGD in MAPS [5] since our previous analysis, we re-evaluated the ancient WGDs with two new MAPS analyses and new data in the 1KP capstone (One Thousand Plant Transcriptomes Initiative, 2019). In one of the MAPS analyses (Fig. 5a), we selected species of *Cicerbita, Lactuca, Tragopogon*, *Leontopodium*, and *Carthamus* to represent the Compositae. Data from *Scaevola* and *Menyanthes* were used as outgroups. Our new analyses with the 1KP data confirmed the phylogenetic position of the paleohexaploidy in the ancestry of the Compositae (Fig. 5a). In the second analysis (Fig. 5b), we used the expanded phylogenetic sampling of the 1KP to more precisely locate an additional WGD in the ancestry of the Heliantheae previously inferred by Ks plots and ortholog divergence analyses [14] and synteny [40]. We selected species of *Flaveria, Xanthium*, and *Helenium* to represent the tribe Heliantheae, and species of *Inula* and four other genera as outgroups. Our analysis of new 1KP data confirmed the location of the Heliantheae WGD with a significant peak of gene duplication consistent with a simulated WGD in the history of all Heliantheae sampled (Fig. 5). Our Compositae analyses in the 1KP allowed us to re-evaluate established WGDs using data from newly sampled taxa and more precisely locate these in the phylogeny. More than 100 of the 1KP WGDs were previously inferred and the expanded sampling of the 1KP dataset allowed us to more precisely place them as we did here in the Compositae.

## 3. Evaluation of WGD inferences

To evaluate our WGD inferences from the 1KP capstone project [1], we compared the consistency of our inferences with whole genome synteny analyses. Although limited in placing WGDs on a phylogeny because of the relatively low phylogenetic sampling of assembled genomes, synteny analysis using high quality genomes is generally considered the best approach for confirming an ancient WGD [37,42]. We compared the results of our Ks and MAPS analyses with analyses of WGDs from published synteny analyses of plant genomes (Fig. 6, SI_Table_1). Overall, we were able to make 65 comparisons of our Ks plot inferences and 43 comparisons of our MAPS phylogenomic analyses to syntenic analyses. Our inferences of WGDs with Ks plots and ortholog divergences were 100% consistent with syntenic analyses from either the same species or a close relative (Fig. 6, SI_Table_1). Despite a perception that Ks plots are difficult to interpret or unreliable, a recent study found that Ks plots analyses using best practices, as we did here, are highly robust [10]. Thus, the high consistency of our Ks plot inferences of WGDs with published genome analyses is not unexpected. We observed slightly lower consistency of our MAPS phylogenomic inferences of WGDs. Across the 43 synteny comparisons, we observed no false positives, but did observe six false negative results (Fig. 6, SI_Table_1). This tendency of our phylogenomic method towards false negatives and “missing” established WGDs is a known issue. There are cases of well established WGDs going undetected with different phylogenomic analyses, including *At*-▯ [43] and the eudicot gamma hexaploidy [31]. Although MAPS and other phylogenomic approaches are often viewed as more rigorous than single species approaches like Ks plots and synteny, these approaches are sensitive to a variety of parameters including gene tree sample size, taxon composition, gene tree occupancy, variation in branch lengths, variation in gene birth/death rates, and variation in gene retention and loss patterns across the phylogeny, to name a few. Notably, we did not observe false positive inferences of WGDs with MAPS, and there does not appear to be reports of false positive inferences in the literature from other phylogenomic methods. However, false signals of large bursts of gene duplication, potentially on the scale consistent with a WGD, could be created by incomplete lineage sorting and quirks of gene tree reconciliation [26]. To minimize the potential biases of these types of phylogenomic methods in the 1KP capstone project, we aimed to use a total evidence approach that combined inferences across Ks plots, ortholog divergence analyses, and MAPS phylogenomic analyses to infer WGDs. Considering that we observed no false positives and high consistency of our Ks and MAPS analyses with syntenic results, we think our survey of WGDs across the phylogeny of green plants is reasonably robust and the combined approach minimized false positives. We expect that some of the 244 WGDs we inferred may move location or be merged as more data become available, and emphasize that the 138 newly inferred WGDs should treated as hypotheses until demonstrated with further data to corroborate the nature and precise timing of these large scale gene duplications.

**Fig. 6.**
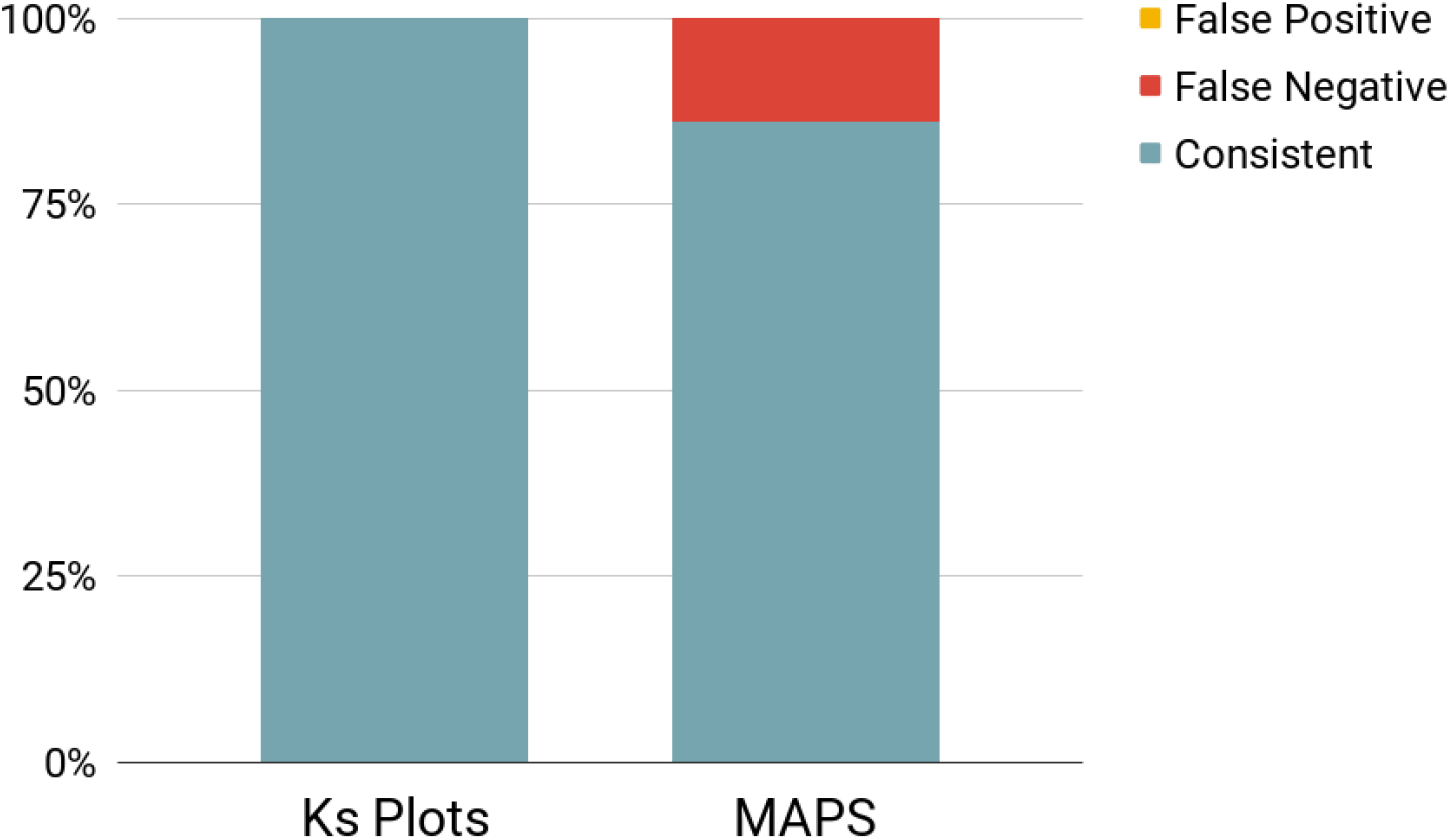
Consistency of the 1KP Ks and MAPS inferences of WGD with results from published synteny analyses of plant genomes. Consistent results represented by blue and false negative results represented by red. There were no false positives in our inferences of WGDs compared to those from published synteny analyses.

## Availability of Supporting Data

https://bitbucket.org/barkerlab/1kp/src/master/

## References

1. One Thousand Plant Transcriptomes Initiative. A phylogenomic view of evolutionary complexity across green plants. Nature.

2. Barker MS, Husband BC, Pires JC. Spreading Winge and flying high: The evolutionary importance of polyploidy after a century of study. Am J Bot. 2016;103:1139–45.

3. Wendel JF. The wondrous cycles of polyploidy in plants. Am J Bot. 2015;102:1753–6.

4. Barker MS, Dlugosch KM, Dinh L, Challa RS, Kane NC, King MG, et al. EvoPipes.net: Bioinformatic Tools for Ecological and Evolutionary Genomics. Evol Bioinform Online. ncbi.nlm.nih.gov; 2010;6:143–9.

5. Li Z, Tiley GP, Galuska SR, Reardon CR, Kidder TI, Rundell RJ, et al. Multiple large-scale gene and genome duplications during the evolution of hexapods. Proc Natl Acad Sci U S A. 2018;115:4713–8.

6. Li Z, Baniaga AE, Sessa EB, Scascitelli M, Graham SW, Rieseberg LH, et al. Early genome duplications in conifers and other seed plants. Sci Adv. American Association for the Advancement of Science; 2015;1:e1501084.

7. Birney E, Clamp M, Durbin R. GeneWise and Genomewise. Genome Res. 2004;14:988–95.

8. Goodstein DM, Shu S, Howson R, Neupane R, Hayes RD, Fazo J, et al. Phytozome: a comparative platform for green plant genomics. Nucleic Acids Res. 2012;40:D1178–86.

9. Yang Z. PAML 4: phylogenetic analysis by maximum likelihood. Mol Biol Evol. 2007;24:1586–91.

10. Tiley GP, Barker MS, Burleigh JG. Assessing the performance of Ks plots for detecting ancient whole genome duplications. Genome Biol Evol [Internet]. 2018; Available from: http://dx.doi.org/10.1093/gbe/evy200

11. Vanneste K, Van de Peer Y, Maere S. Inference of genome duplications from age distributions revisited. Mol Biol Evol. SMBE; 2013;30:177–90.

12. Cui L, Wall PK, Leebens-Mack JH, Lindsay BG, Soltis DE, Doyle JJ, et al. Widespread genome duplications throughout the history of flowering plants. Genome Res. genome.cshlp.org; 2006;16:738–49.

13. Benaglia T, Chauveau D, Hunter D, Young D. mixtools: An R Package for Analyzing Mixture Models. Journal of Statistical Software, Articles. 2009;32:1–29.

14. Barker MS, Kane NC, Matvienko M, Kozik A, Michelmore RW, Knapp SJ, et al. Multiple paleopolyploidizations during the evolution of the Compositae reveal parallel patterns of duplicate gene retention after millions of years. Mol Biol Evol. 2008;25:2445–55.

15. Shi T, Huang H, Barker MS. Ancient genome duplications during the evolution of kiwifruit (Actinidia) and related Ericales. Ann Bot. 2010;106:497–504.

16. Barker MS, Vogel H, Schranz ME. Paleopolyploidy in the Brassicales: analyses of the Cleome transcriptome elucidate the history of genome duplications in Arabidopsis and other Brassicales. Genome Biol Evol. 2009;1:391–9.

17. Cannon SB, McKain MR, Harkess A, Nelson MN, Dash S, Deyholos MK, et al. Multiple polyploidy events in the early radiation of nodulating and nonnodulating legumes. Mol Biol Evol. 2015;32:193–210.

18. Yang Y, Moore MJ, Brockington SF, Soltis DE, Wong GK-S, Carpenter EJ, et al. Dissecting Molecular Evolution in the Highly Diverse Plant Clade Caryophyllales Using Transcriptome Sequencing. Mol Biol Evol. 2015;32:2001–14.

19. Roodt D, Lohaus R, Sterck L, Swanepoel RL, Van de Peer Y, Mizrachi E. Evidence for an ancient whole genome duplication in the cycad lineage. PLoS One. 2017;12:e0184454.

20. Smith SA, Brown JW, Yang Y, Bruenn R, Drummond CP, Brockington SF, et al. Disparity, diversity, and duplications in the Caryophyllales. New Phytol. 2018;217:836–54.

21. Emms DM, Kelly S. OrthoFinder: solving fundamental biases in whole genome comparisons dramatically improves orthogroup inference accuracy. Genome Biol. 2015;16:157.

22. Mirarab S, Nguyen N, Warnow T. PASTA: Ultra-Large Multiple Sequence Alignment. In: Sharan R, editor. Research in Computational Molecular Biology. Cham: Springer International Publishing; 2014. p. 177–91.

23. Katoh K, Misawa K, Kuma K-I, Miyata T. MAFFT: a novel method for rapid multiple sequence alignment based on fast Fourier transform. Nucleic Acids Res. 2002;30:3059–66.

24. Edgar RC. MUSCLE: multiple sequence alignment with high accuracy and high throughput. Nucleic Acids Res. 2004;32:1792–7.

25. Stamatakis A. RAxML version 8: a tool for phylogenetic analysis and post-analysis of large phylogenies. Bioinformatics. 2014;30:1312–3.

26. Hahn MW. Bias in phylogenetic tree reconciliation methods: implications for vertebrate genome evolution. Genome Biol. 2007;8:R141.

27. Smith SA, Moore MJ, Brown JW, Yang Y. Analysis of phylogenomic datasets reveals conflict, concordance, and gene duplications with examples from animals and plants. BMC Evol Biol. 2015;15:150.

28. Rabier C-E, Ta T, Ané C. Detecting and locating whole genome duplications on a phylogeny: a probabilistic approach. Mol Biol Evol. 2014;31:750–62.

29. Sjöstrand J, Arvestad L, Lagergren J, Sennblad B. GenPhyloData: realistic simulation of gene family evolution. BMC Bioinformatics. 2013;14:209.

30. Jiao Y, Wickett NJ, Ayyampalayam S, Chanderbali AS, Landherr L, Ralph PE, et al. Ancestral polyploidy in seed plants and angiosperms. Nature. 2011;473:97–100.

31. Zwaenepoel A, Van de Peer Y. Inference of Ancient Whole-Genome Duplications and the Evolution of Gene Duplication and Loss Rates. Mol Biol Evol. 2019;36:1384–404.

32. Ruprecht C, Lohaus R, Vanneste K, Mutwil M, Nikoloski Z, Van de Peer Y, et al. Revisiting ancestral polyploidy in plants. Sci Adv. 2017;3:e1603195.

33. Barker MS, Li Z, Kidder TI, Reardon CR. Most Compositae (Asteraceae) are descendants of a paleohexaploid and all share a paleotetraploid ancestor with the Calyceraceae. American Journal of [Internet]. Wiley Online Library; 2016; Available from: https://onlinelibrary.wiley.com/doi/abs/10.3732/ajb.1600113

34. Huang C-H, Zhang C, Liu M, Hu Y, Gao T, Qi J, et al. Multiple Polyploidization Events across Asteraceae with Two Nested Events in the Early History Revealed by Nuclear Phylogenomics. Mol Biol Evol. SMBE; 2016;33:2820–35.

35. Jiao Y, Leebens-Mack J, Ayyampalayam S, Bowers JE, McKain MR, McNeal J, et al. A genome triplication associated with early diversification of the core eudicots. Genome Biol. 2012;13:R3.

36. Jaillon O, Aury J-M, Noel B, Policriti A, Clepet C, Casagrande A, et al. The grapevine genome sequence suggests ancestral hexaploidization in major angiosperm phyla. Nature. 2007;449:463–7.

37. Lyons E, Pedersen B, Kane J, Alam M, Ming R, Tang H, et al. Finding and comparing syntenic regions among Arabidopsis and the outgroups papaya, poplar, and grape: CoGe with rosids. Plant Physiol. 2008;148:1772–81.

38. Vekemans D, Proost S, Vanneste K, Coenen H, Viaene T, Ruelens P, et al. Gamma paleohexaploidy in the stem lineage of core eudicots: significance for MADS-box gene and species diversification. Mol Biol Evol. 2012;29:3793–806.

39. Reyes-Chin-Wo S, Wang Z, Yang X, Kozik A, Arikit S, Song C, et al. Genome assembly with in vitro proximity ligation data and whole-genome triplication in lettuce. Nat Commun. 2017;8:14953.

40. Badouin H, Gouzy J, Grassa CJ, Murat F, Staton SE, Cottret L, et al. The sunflower genome provides insights into oil metabolism, flowering and Asterid evolution. Nature. 2017;546:148–52.

41. Song C, Liu Y, Song A, Dong G, Zhao H, Sun W, et al. The Chrysanthemum nankingense Genome Provides Insights into the Evolution and Diversification of Chrysanthemum Flowers and Medicinal Traits [Internet]. Molecular Plant. 2018. p. 1482–91. Available from: http://dx.doi.org/10.1016/j.molp.2018.10.003

42. Tang H, Bowers JE, Wang X, Ming R, Alam M, Paterson AH. Synteny and Collinearity in Plant Genomes [Internet]. Science. 2008. p. 486–8. Available from: http://dx.doi.org/10.1126/science.1153917

43. Tiley GP, Ané C, Burleigh JG. Evaluating and Characterizing Ancient Whole-Genome Duplications in Plants with Gene Count Data. Genome Biol Evol. 2016;8:1023–37.

